# Crystal structure of Senecavirus A 3C protease

**DOI:** 10.1101/2020.02.21.959106

**Authors:** Kaiwen Meng, Lijie Zhang, Geng Meng

## Abstract

Senecavirus A (SVA), an emerging picornavirus in porcine population, could infect porcines of all age group and cause FMD-like symptoms. Picornaviridae, a group of RNA viruses do harm to both human and stocks; however, most of picornaviruses are lack of effective vaccines and drugs. Picornaviral 3C protease (3C^pro^), as an important role in virus maturation, they basically take charge of poly-protein cleavaging, RNA replication, and multiple interventions on host cells. In this study, we successfully solved the crystal structure of 3C^pro^ at 1.9 Å resolution. The results showed several differences of the binding groove within picornaviral 3C^pro^, and prompted that the accommodate ability of the pocket may associate with the cleavage efficiency. The further research on 3C^pro^ cleavage efficiency based on structural biology, will prospectively provide an instruction on designing of efficient 3C^pro^ for universally proteolysis in picornaviral VLP production.

## Introduction

Senecavirus A (SVA), once called Seneca Valley Virus (SVV), is an emerging virus in porcine population that has been popular worldwide since 2015. SVA could infect porcine of all age group, clinically manifest symptoms indistinguishable from those of foot-and-mouth disease (FMD) (Zhang et al., 2018).

SVA belongs to the genus *Senecavirus*, family *Picornaviridae*. The virus genome consists of a positive sense, single-stranded RNA of ∼7280nt in length, encoding a large precursor poly-protein. 3C protease (3C^pro^), a chymotrypsin-like cysteine protease shared by all Picornavirus, plays an important role in virus maturation, they basically take charge of poly-protein’s proteolytic processing. Ten of the thirteen cleavage sites of the poly-protein are carried out by the 3C protease (3C^pro^), including VP2 / VP3, VP3 / VP1, VP1 / 2A, 2B / 2C, 2C / 3A, 3A / 3B1, 3B1 / 3B2, 3B2 / 3B3, 3B3 / 3C and 3C / 3D. According to Berger and Schechter’s nomenclature (Berger and Schechter, 1970), the residues within the substrate preceding and following the cleavage site is denoted P and P’, respectively; and those subsites within the 3C^pro^ that accommodate P or P’ residues are numbered as S and S’. Most picornaviral 3C^pro^ exhibit marked preference for P1, and P1’ residue types (Gln–Gly junctions), whereas the rest of the P and P’ positions show less sequence conservation. Surprisingly, mutations at those less conservative positions would dramatically influence the rate of peptide cleavage by 3C^pro^ (Lu et al., 2011).

As Picornavirus basically rely on 3C^pro^ to generate the individual structural and nonstructural proteins, 3C^pro^ is a necessary component for Picornaviral virus-like particle (VLP) production. However, the limitation of the use of 3C^pro^ is considerably significant: the tolerance of expression system against protease activity, the proportion between structural protein and 3C^pro^ expression, and the cleavage efficiency issue (Belsham and Bøtner, 2015; Polacek et al., 2013; Porta et al., 2013).

There are already a lot of work reveals the structural basis for 3C^pro^’s substrate recognition, as well as the relationship between proteolytic efficiency and cleavage site mutations. We believed that the structural information of 3C^pro^ will offer an instruction to us to design efficient 3C^pro^ for universally proteolysis in picornaviral VLP production.

## Results

### Phylogenetics and cleavage specificities of SVA 3C^pro^

Phylogenetic comparison of the sequence of SVA 3C^pro^ with those of other picornaviruses suggests that SVA is related closely to Encephalomyocarditis virus (EMCV) and Human TMEV-like cardiovirus (HTCV), and more closely to Foot-and-mouth disease virus (FMDV) among all structure solved picornaviral 3C^pro^ (Fig.1).

**Figure 1.**
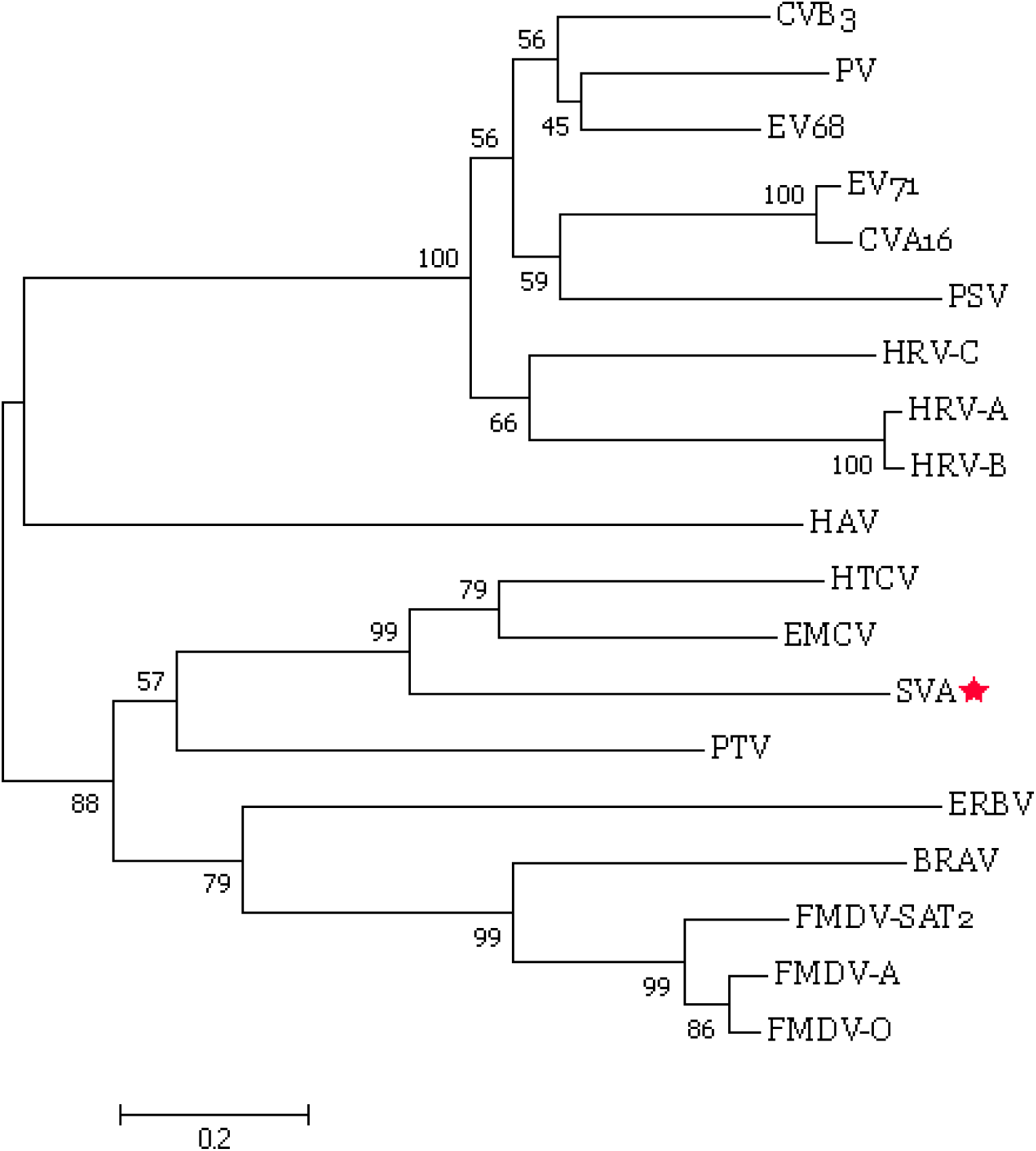
Phylogenetic comparison of the picrnaviral 3C^pro^ sequences. Midpoint-rooted Neighbor-joining tree were produced in MEGA 7 (Kumar et al., 2016) using the Poisson model. The number of bootstrap replicates were set as 2000. Bootstrap values are given at the nodes.

We carried out an investigation on the sequence variation of the cleavage site associate with structural proteins (Fig. 2). The four distinct structural proteins of SVA are termed VP1-4, which are responsible for inducing humoral and cellular immunity in the animal body. As other picornavirus, the SVA 3C^pro^ can specifically cleave P1 region into VP0, VP1 and VP3. VP0 will be further cleavaged into VP2 and VP4 during the encapsidation of RNA by unknown mechanism(Belsham and Bøtner, 2015). The cleavage site of SVA between VP0-VP3 is Q/G, which is consistent with the cleavage site of most picornaviruses. Whereas the cleavage site between VP1-VP3 is H/S, which is uncommon.

**Figure 2.**
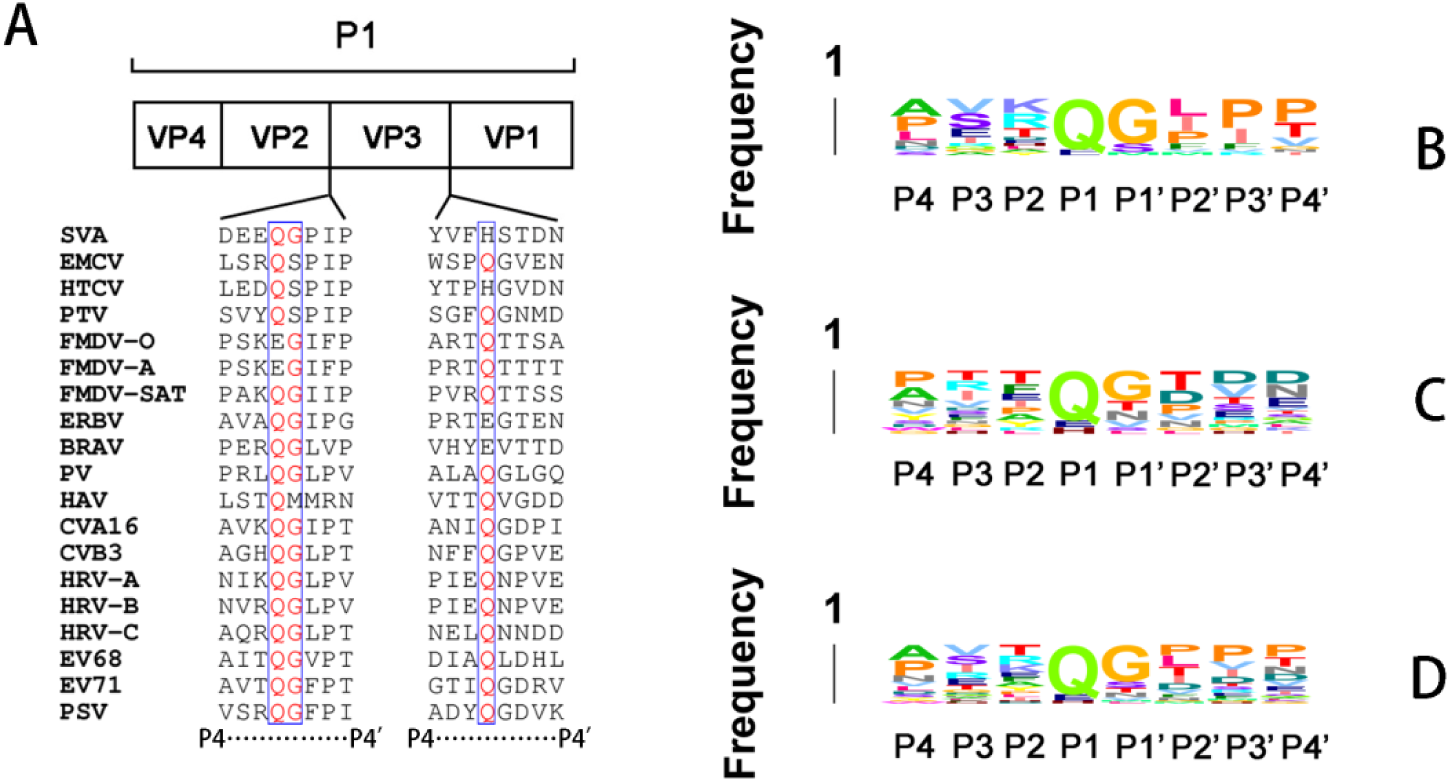
(A) Overview of P1 region organization within the picornaviral polyproteins. The two cleavage junction sites that are to be processed by 3C^pro^ are indicated, with each joining sequence spanning from the P4 to P4’ residues listed below the schematic. (B) Sequence logo of VP0-VP3 cleavage site. (C) Sequence logo of VP3-VP1 cleavage site. (D) Sequence logo of both cleavage sites.

Alignment of peptides spanning four residues either side corresponding to VP0-VP3 and VP1-VP3 cleavage positions, reveals that picornaviral 3C^pro^ share similar substrate specificity in recognizing small hydrophobic amino acid residues at most positions, except P1 and P2. The P1 position is invariably occupied by hydrophilic amino acid, and in most cases it is occupied by a glutamine. On the contrary, P2 position shows no significant preference on amino acid residues, whether it is acidic, basic, aliphatic or polar.

However, the preference for small hydrophobic residues on most position is not an absolute. For SVA, the P4 position in VP0-VP3 and VP1-VP3 cleavage sequence were strikingly occupied by bigger amino acid residues, aspartic acid and tyrosine respectively, instead of alanine and proline in most cases. For other picornaviruses, seen in Fig. 2, amino acid residue with bigger side chain and/or different property also can be accommodate by a same subsite, that makes the 3C^pro^ possible to recognize and cleave different site within the poly-protein. But this tolerance also brings notable variety in cleavage efficiency(Birtley et al., 2005; Lu et al., 2011; Zunszain et al., 2010). This phenomenon can be explained by the subsites’ structure and will be discussed later.

### Overall structure of SVA 3C^pro^

The 3C^pro^ exists as a monomer in solution, as indicated by gel filtration (not shown). Crystals grew from 0.18 M lithium chloride (pH 7.0), 12 to 18% PEG 3350. The crystal structure of SVA 3C^pro^ was determined at 1.9 Å resolution by the molecular replacement. The final refinement of the structure generated the R/Rfree factors of 18.43/23.55% (Table 1). The crystal belongs to space group P 1 2_1_ 1, with two 3C^pro^ monomers per asymmetric unit. The two molecules are very similar, and the root mean square deviation (RMSD) for all of the Cα atoms in two molecules was 0.258 Å. Therefore, unless otherwise noted, in comparisons below we shall be referring to one molecule in the asymmetric unit.

**Table 1.**
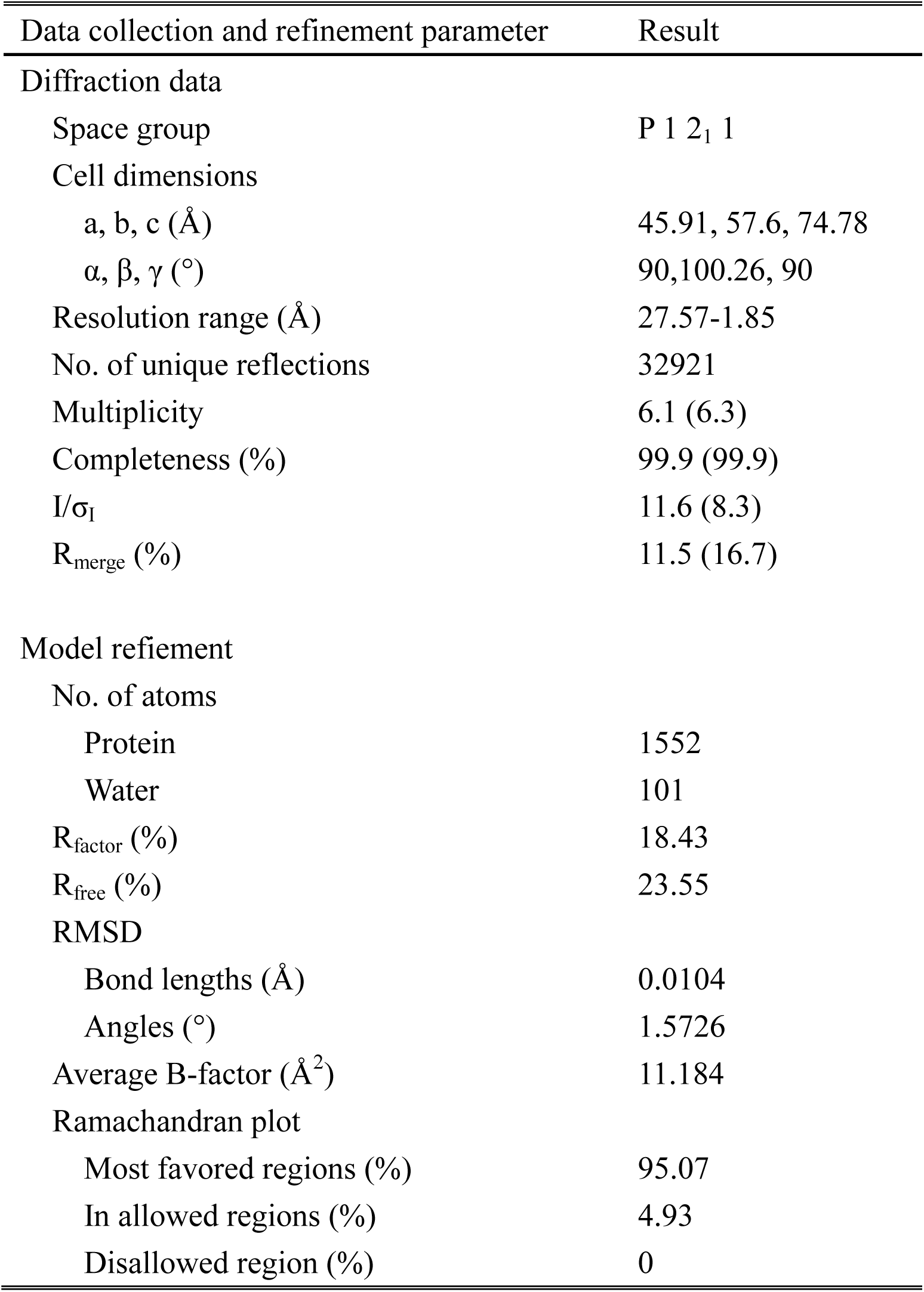
Data collection and refinement statistics.

| Data collection and refinement parameter | Result |
| --- | --- |
| Diffraction data |  |
| Space group | P 1 2<sub>1</sub> 1 |
| Cell dimensions |  |
| a, b, c (Å) | 45.91, 57.6, 74.78 |
| $\alpha$, $\beta$, $\gamma$ (°) | 90, 100.26, 90 |
| Resolution range (Å) | 27.57-1.85 |
| No. of unique reflections | 32921 |
| Multiplicity | 6.1 (6.3) |
| Completeness (%) | 99.9 (99.9) |
| I/σ<sub>I</sub> | 11.6 (8.3) |
| R<sub>merge</sub> (%) | 11.5 (16.7) |
| Model refinement |  |
| No. of atoms |  |
| Protein | 1552 |
| Water | 101 |
| R<sub>factor</sub> (%) | 18.43 |
| R<sub>free</sub> (%) | 23.55 |
| RMSD |  |
| Bond lengths (Å) | 0.0104 |
| Angles (°) | 1.5726 |
| Average B-factor (Å<sup>2</sup>) | 11.184 |
| Ramachandran plot |  |
| Most favored regions (%) | 95.07 |
| In allowed regions (%) | 4.93 |
| Disallowed region (%) | 0 |

SVA 3C^pro^ adopts a typical chymotrypsin fold that is similar to those of other picornaviral 3C^pro^. The overall structure of SVA 3C^pro^ is shown in Fig. 3. It contains 204 amino acids from residues D3 to R206, forming two domains. The first domain is largely composed of a 7-stranded β-barrel structure (A1 to G1). The second domain also contains a compact barrel core, which is composed of 8 β-strands (A2 to H2) arranged in an antiparallel manner. Overall, the two domains are connected via a long loop (amino acids 90 to 113 aa) over the “rear” surface of the molecule. The catalytic triad of His 48, Asn 84, Cys 160 is located in the cleft formed by the two β-barrel domains.

**Figure 3.**
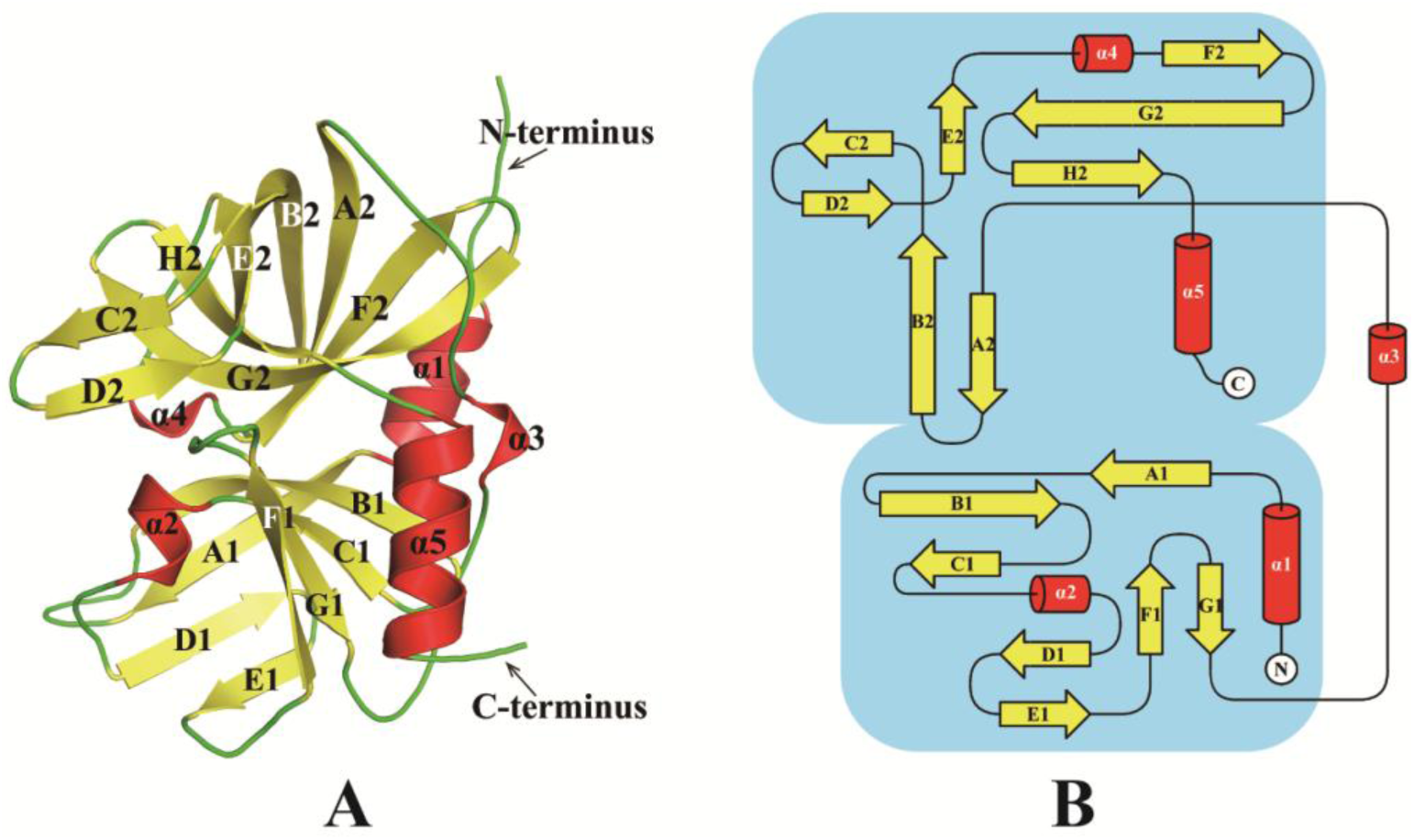
The overall structure of SVA 3C^pro^. (A) Cartoon representation of the structure of SVA 3C^pro^. (B) Topology diagram of SVA 3C^pro^.

### Comparison of SVA 3C^pro^ and related picornaviral protease

Although they belong to the same family, the multi-sequences alignment shows that SVA 3C^pro^ only shares ≤40 % amino acid sequence identity with other picornaviral 3C^pro^ (Fig. 4), among which 3C^pro^ of EMCV shares the highest sequence similarity of 37.36%, while EV71 3C^pro^ shares the lowest similarity, 15.8%. In addition, FMDV Type A 3C^pro^ also shows as high as 23.86% sequence align percent identity with SVA 3C^pro^, which is the highest among all structure solved picornaviral 3C^pro^. Only three amino acid residues are invariable among all reference sequences, including H48, G158 and G161. H48 is one of the catalytic residue, and the latter two are associated with catalytic important motif G-X-C/S-G-G. Besides, G176, H178, G181 located on G2 strand are also highly conserved. While SVA 3C^pro^ possess a relatively conservative catalytic motif as other picornaviral 3C^pro^, it doesn’t have a characteristic KFRDI motif. The KFRDI motif was previously characterized as one of RNA binding motifs in picornaviral 3C^pro^, it is mutated into SFPNN (95-99aa) in SVA 3C^pro^(Leong et al., 1993; Matthews et al., 1994; Mosimann et al., 1997; Walker et al., 1995).

**Figure 4.**
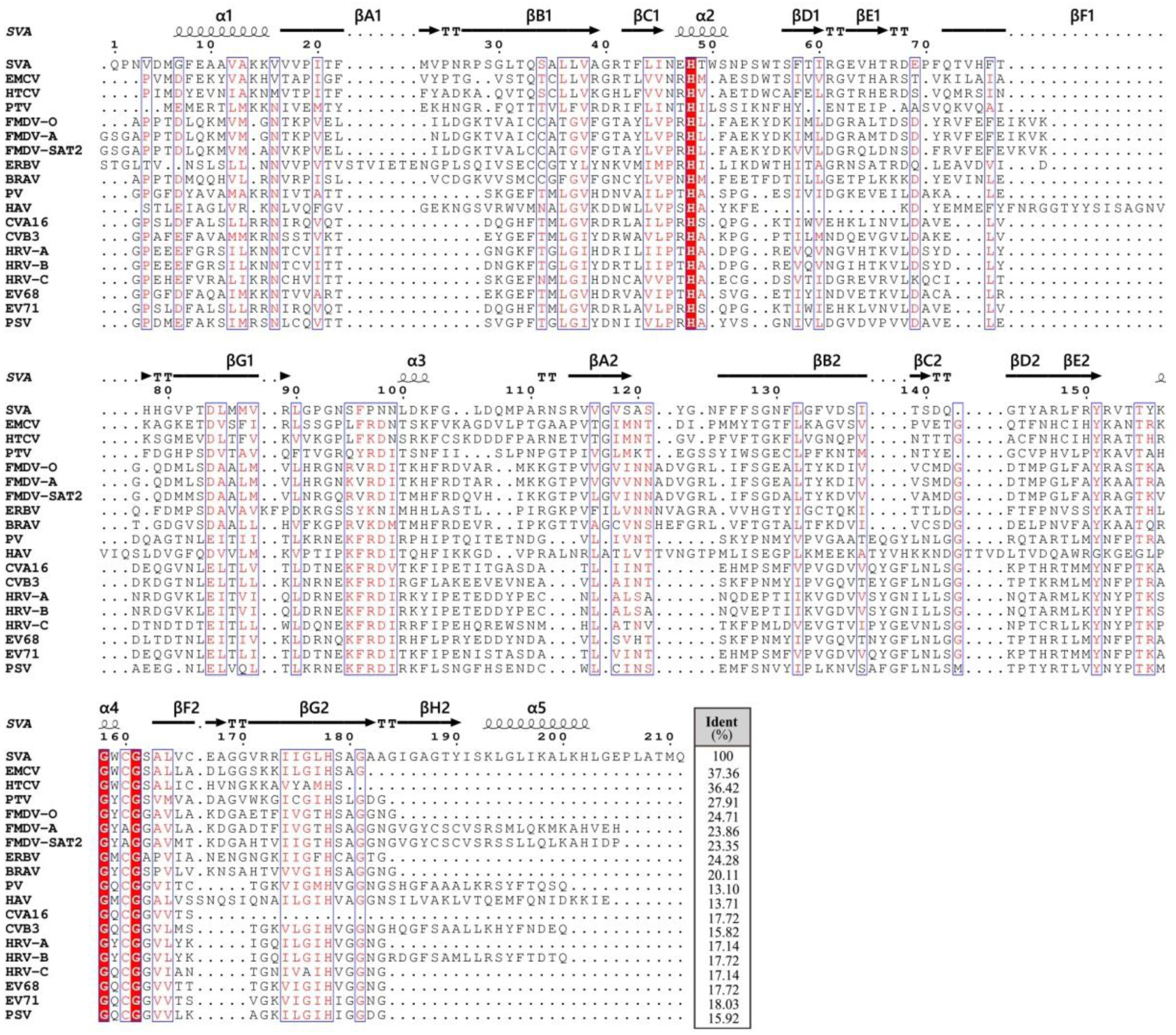
Structure based multiple-sequence alignment of the 3C^pro^ from common picornaviruses. Invariant residues in 3C pro are highlighted with red background; conserved residues are shown in red font.

Overall, despite SVA 3C^pro^ shares low homology with other picornaviral 3C^pro^, SVA 3C^pro^ still maintains the classic chymotrypsin fold. Superposition of the SVA 3C^pro^ structure with the other solved picornaviral 3C^pro^ structures resulted in a root mean square difference (RMSD) in α-carbon positions about 1.6-1.9 Å (Table 2). Here we focus on five positions with sequence insertion, which resulting in structural differences in SVA 3C^pro^(Fig. 5).

**Table 2.** Alignment of SVA 3Cpro and other known 3C^pro^s.

|  | N-terminus<br>RMSD(Å) | Loop<br>RMSD(Å) | C-terminus<br>RMSD(Å) | Overall<br>RMSD(Å) |
| --- | --- | --- | --- | --- |
| Without substrate |  |  |  |  |
| SV-PV (1L1N) | 1.823 | 2.623 | 1.465 | 1.784 |
| SVA-CVA16 (3SJ8) | 1.641 | 2.582 | 1.492 | 1.703 |
| SVA-CVB3 (2ZU1) | 2.135 | 2.160 | 1.654 | 1.927 |
| SVA-EV68 (3ZV8) | 2.024 | 2.254 | 1.635 | 1.876 |
| SVA-EV71 (3OSY) | 1.683 | 2.514 | 1.682 | 1.798 |
| SVA-FMDV A10 (2BHG) | 1.611 | 1.958 | 1.250 | 1.515 |
| SVA-FMDV SAT2 (5HM2) | 1.657 | 2.338 | 1.335 | 1.607 |
| SVA-HAV (1HAV) | 2.180 | 2.019 | 1.689 | 1.944 |
| With substrate |  |  |  |  |
| SVA-PV (4DCD) | 1.759 | 2.200 | 1.445 | 1.680 |
| SVA- CVA16 (3SJ9) | 1.731 | 2.654 | 1.466 | 1.747 |
| SVA-CVB3 (2ZU3) | 2.029 | 2.093 | 1.599 | 1.829 |
| SVA-EV68 (3ZVA) | 2.064 | 2.236 | 1.502 | 1.893 |
| SVA-EV71 (3SJO) | 1.682 | 2.568 | 1.509 | 1.731 |
| SVA-HRV2 (1CQQ) | 1.855 | 2.415 | 1.495 | 1.768 |
| SVA-FMDV A10 (2WV5) | 1.568 | 1.973 | 1.100 | 1.424 |
| SVA- HAV(2HAL) | 2.163 | 2.037 | 1.386 | 1.828 |

**Figure 5.**
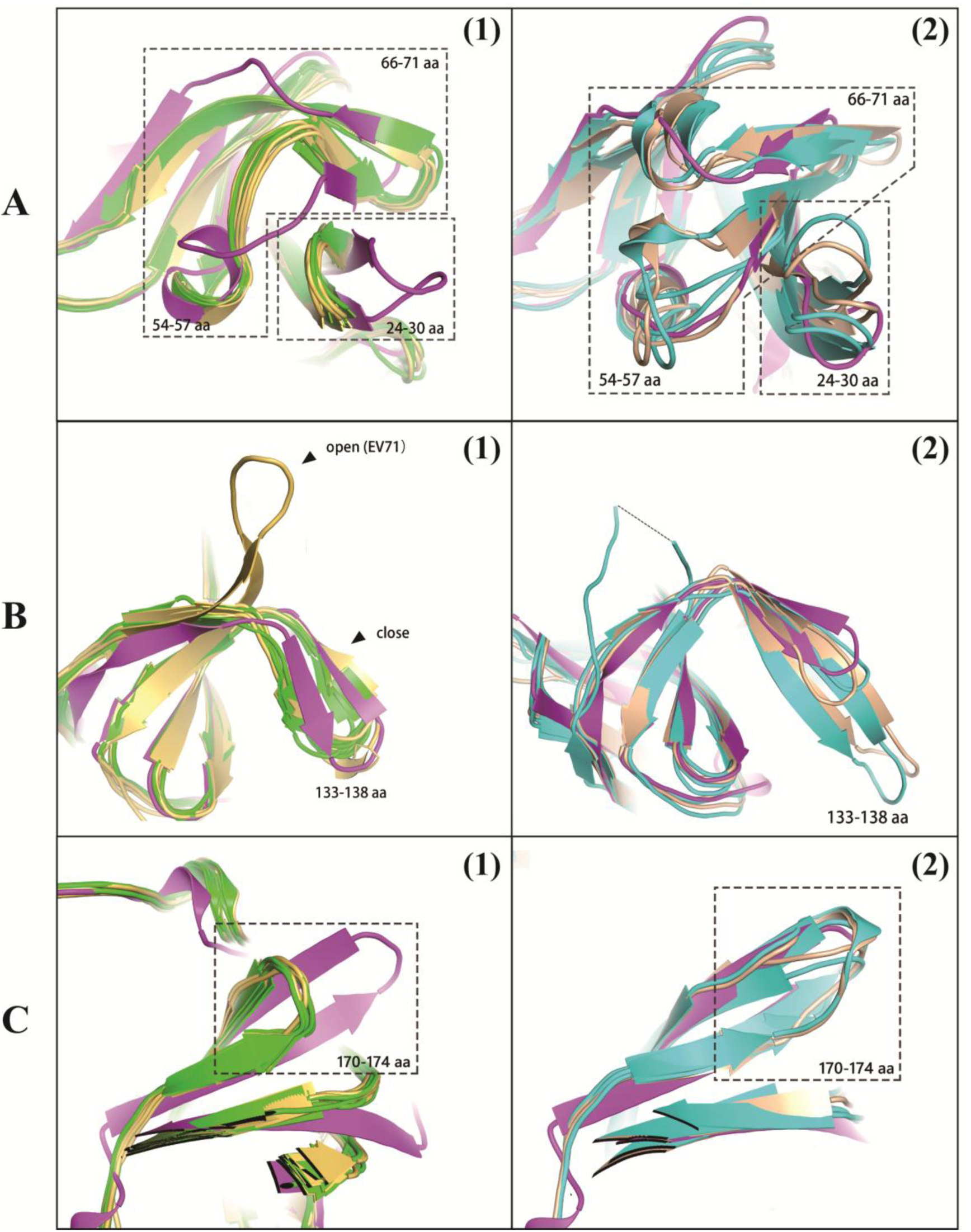
Superposition of SVA 3C^pro^(magneta) on other known 3C^pro^s. The superposition is separated into two groups, in order to exhibt major differences legibly. Group 1 (without substrate – yellow, with substrate – green) including enterovirus and rhinovirus; group 2 (without substrate – cyan, with substrate – wheat) including FMDV and HAV. (A) Cartoon representation of major difference within 24-30aa, 54-57aa and 66-71 aa. (B) Cartoon representation of major difference within 133-138 aa. (C) Cartoon representation of major difference within 170-174 aa.

The first sequence insertion occurs between 24-30aa. With six more amino acid residues, SVA 3C^pro^ contains a longer loop between βA1-B1 strands further away from the substrate binding groove than PV(1L1N, 4DCD), CVA16(3SJ8, 3SJ9), CVB3(2ZU1, 2ZU3), EV68(3ZV8, 3ZVA) and EV71 (3OSY, 3SJO). The superposition consequence shows that short loop between βA1-βB1 strands will only adopt a slight change on angle after substrate combination. On the contrast, the longer loops of the 3C^pro^ from FMDV-A(2BHG, 2WV5), FMDV-SAT2(5HM2) and HAV(1HAV, 2HAL) resulting from sequence insertion shows significant transformation after combining substrate. In FMDV 3C^pro^-peptide complex, the loop between βA1-B1 strands bend away from the cleft; and in HAV 3C^pro^ -inhibitor complex, the loop move toward the binding groove. Considering about 24-30aa are associate with the recognition of the P’ portion of substrate, these result implies that the βA1-B1 strands of SVA 3C^pro^ might be a flexible structure.

Locate into the structure, the α2 helix to βF1 strand segment possibly participates in forming the side wall and floor of the S’ pocket. Residues insertion between 54-57aa and 66-71aa in SVA 3C^pro^ resulting in a longer α2 helix, which draws the loop between α2 helix and βD1 strand near to βA1-B1 strands, meanwhile βD1-E1 strands bends away from the protein surface. The continuous βE1 strand in the 3C^pro^ from PV, CVA, CVB, EV68 and EV71 is broken into βE1 and βF1 in SVA, FMDV and HAV. A short loop protruding from the outer surface is formed between βE1 and βF1 in SVA 3C^pro^, while in the 3C^pro^ from FMDV and HAV a short helix located in that position. In the α2 helix to βE1 strand segments, SVA 3C^pro^ shares similar structure with FMDV 3C^pro^. However, HAV 3C^pro^ is less structure conserved between α2 helix and βE1 strand when compared with SVA 3C^pro^ and FMDV 3C^pro^: a shorter α2 helix followed by a loop with similar orientation of PV, CVA, CVB, EV68 and EV71; an extra helix between α2 helix and βD1 strand; and a longer βD1-E1 strand.

The flexible surface loop between βC2 and βD2 strands, denoted as β-ribbon, whose configuration is not affected by the sequence insertion between 133-138aa. The β-ribbon plays an important role in recognizing the P2–P4 region of peptide substrates by transforming between two conformations (“open” and “close”), the mobility of the β-ribbon is relevant to protease activity(Cui et al., 2011). SVA 3C^pro^ in this study retain this β-ribbon in a close state, i.e. the loop is located over the substrate binding groove with its apical tip pointing toward the protease active site.

The last sequence insertion is observed between 170-174aa. Thus, SVA 3C^pro^ has a longer βF2-G2 strands orient differ from PV, CVA, CVB, EV68 and EV71. In PV, CVA, CVB, EV68 and EV71, βF2-G2 strands in 3C^pro^ bent toward α1 helix. The FMDV and HAV shares similar βF2-G2 strand with SVA 3C^pro^, albeit a short helix was formed on the tip in HAV 3C^pro^.

In general, both sequence and structural alignments indicate that SVA 3C^pro^ has a higher variability in the N-terminal domain compared with other known picornaviral 3C^pro^, while the C-terminus is relatively conservative. Furthermore, piles of researches were focused on C terminus and revealed that the C-terminus plays an important role in substrate recognition.

### Prediction of the subsite conformation in SVA 3C^pro^

In order to identify residues of SVA 3C^pro^ directly involve in substrate binding, we superimpose the SVA 3C^pro^ onto the FMDV 3C^pro^-APAKELLNF peptide co-crystal structure, CVA 3C^pro^-GLRQAVTQ peptide co-crystal structure and HRV 3C^pro^-AG7088 co-crystal structure (Fig.6). The width of S subsite within SVA 3C^pro^ and FMDV 3C^pro^ are shown in table 3.

**Figure 6.**
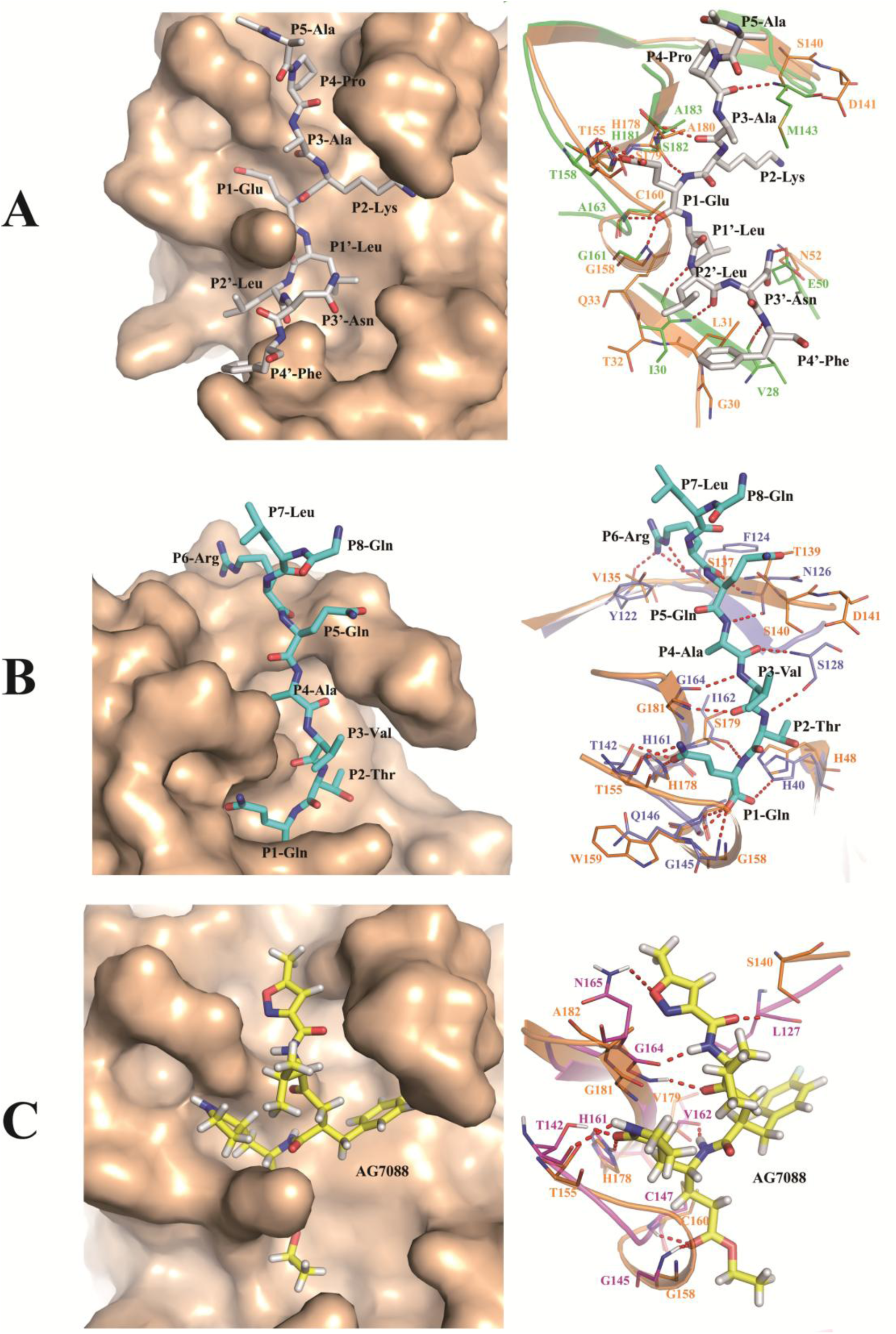
Prediction on residues of SVA 3Cpro directly involve in substrate binding. (A) Superimpose the SVA 3C^pro^ (orange) onto the FMDV 3Cpro-APAKELLNF peptide co-crystal structure. The peptide is shown as white, and the residues interact with the substrate is shown as green lines. (B) Superimpose the SVA 3C^pro^ onto the CVA 3Cpro-GLRQAVTQ peptide co-crystal structure. The peptide is shown as cyan, and the residues interact with the substrate is shown as light blue lines. (C) Superimpose the SVA 3C^pro^ onto the HRV 3Cpro-AG7088 co-crystal structure. The inhibitor is shown as yellow, and the residues interact with the substrate is shown as magneta lines.

**Table 3.** Width of the S subsite within picornaviral 3C^pro^s.

|  | SVA | FMDV |
| --- | --- | --- |
| S4 | 11.1 Å | 9.2 Å |
| S3* | NA | NA |
| S2 | 9.1 Å | 9.5 Å |
| S1 | 6.9 Å | 7.8 Å |
| S1' | 10.9 Å | 12 Å |
| S2' | 8.3 Å | 8.6 Å |
| S3' | 7.0 Å | 6.9 Å |
| S4'* | NA | NA |

\* S3 and S4' shows no significant pocket structure.

In picornavirus, P4 residue exhibits conservatism, whose favored amino acid is valine, proline and alanine. However, as mentioned before, the P4 position in VP0-VP3 and VP1-VP3 cleavage sequence of SVA are occupied by amino acid residues with larger side chain, i.e. aspartic acid and tyrosine. This phenomenon can be explained by the conformation of the S4 subsite. In SVA 3C^pro^, the S4 subsite pocket wall is formed by 135-141aa and 182-186aa. Among them, T139, S140, D141 and A182 may interact with P4 residue according to the superposition result. It is noteworthy that in SVA 3C^pro^, these positions are variable and further away from the center of the substrate binding groove than the other structures in the reference. Thus, the wider S4 subsite might be a clue that residues larger than valine or proline can be tolerate by the protease. As for the tolerance of lager side chain in limited S4 subsite, e.g. the S4 position of HRV is limited but able to accommodate an asparagine (Fig. 2A), also can be explained by these interacted positions: an H-bond formed by peptide backbone atoms only, suggesting a degree of side-chain variation at these positions is acceptable.

Many structure solved picornaviral 3C^pro^ haven’t shown significant S3 pocket(Curry et al., 2007; Zunszain et al., 2010). The superimpose result indicate A180 or G181 within SVA 3C^pro^ may contact P3 with backbone H-bond. And this positions are highly conserved in picornavirus.

As previously noted, the P2 residues of the picornaviral substrate peptide are variable. The S2 subsite is surrounded by β-ribbon, α2 helix and the loop between βF1 and βG1 strands, which is a pocket with enough size to accommodate various residues. However, it is seemed that the superposition doesn’t provide much clue on the direct interaction between P2 and S2. In the reference structure, only CVA 3C^pro^ shows one interaction position, one backbone H-bond formed between S128 and P2. But in SVA 3C^pro^, the corresponding position is too far to interact with P2.

The P1 residue can be superimposed into the pocket formed by 155-160aa and 178-180aa of SVA 3C^pro^. H48 within α2 helix, T155, G158, W159, C160, H178 and S179 are seemed possible to form H-bonds with both main chain and side chain of P1 residue. The above positions are all conserved among picornaviral 3C^pro^, except for S179. In Enterovirus, it is a hydrophobic amino acid residue valine or isoleucine interact with P1 backbone instead of a serine with hydrophilicity. Moreover, the difference within 158-162aa is notable. 159-162aa is corresponding to the G-X-C/S-G-G motif, which is highly conserved in picornaviral 3C^pro^. In SVA 3C^pro^, the G162 is mutated into serine, and forms extra H-bonds with S119 from βA2 strand and neighboring W159. What’s more, an α helix was formed with the participation of G158 and W159, which is not observed in other picornaviral 3C^pro^. The extra α helix drags 156-158aa closer to the substrate binding groove. The large side chain from K157 formed an fingerlike projection over S1 and S2 subsites with its tips pointing to the β-ribbon. All these transformation makes the S1 subsite of SVA 3C^pro^ smaller than the other.

There’s not many research concern about the recognition of P’ portion. The superposition of SVA 3C^pro^ onto the FMDV 3C^pro^-APAKELLNF peptide co-crystal structure implies that G30, L31, T32, Q33, N52 might associate with the forming of S’ pockets. According to the measurement result (table 3), SVA S1’ pocket is smaller than FMDV, while S2’ and S3’ are approximately equivalent, suggesting SVA 3C^pro^ may have higher specifity to sequence variance of P’ portion. As for S4’, the subsite that accommodate P4 in FMDV 3C^pro^ are occupied by large side chain in SVA 3C^pro^, which causes in the superimpose result P4 pierces into the surface of SVA 3C^pro^ (Fig. 6A).

## Discussion

We report here the first structural insights into SVA 3C^pro^. Our result reveals several unexpected differences of the structure between SVA 3C^pro^ and other known picornaviral 3C^pro^. The main difference is SVA 3C^pro^ adopts a wider S4 pocket and smaller S1, S1’ pockets.

A previous research on EV71 and CVA16(Lu et al., 2011) suggested that mutate small amino acid residue into larger one at P4 and P1’ positions could dramatically reduce the rate of peptide cleavage. They explained this phenomenon by observing the limited S4 subsite in both EV71 and CVA16 3C^pro^s, and reckoned S1’ subsite as a small subsite too. According to their study, the protease efficiency is restrict by limitied P4 and S1’ pocket when come across with larger amino acid side chain within cleavage sequence. In this regard, we propose a conjecture about the relationship between the structure of the binding groove and the cleavage efficiency. The capacity of the binding groove might be related to the cleavage efficiency. Although small pockets may be able to accommodate large side chain amino acids due to they can only form H- bonds with the peptide backbone, but their cleavage efficiency may still significant reduce when encountering larger side chain amino acids. This influence might be weakened by enlarged pocket or more flexible pocket. The latter situation can be achieved by steric hindrance changes, lower steric hindrance can enhance the flexibility of residues nearby binding pocket, and finally result in the increasing of the pockets’ accommodation ability. It should be noted that too wide binding grooves could also slightly impaired the efficiency probably because of P residue is bound less tightly in the enlarge pocket(Zunszain et al., 2010).

According to Lu G’s paper(Lu et al., 2011), CVA16 3C^pro^ could cleave its own structural protein as well as structural protein of EV71 with surprisingly higher efficiency than EV71. We look back into the structures of EV71 3C^pro^ and CVA16 3C^pro^, the result seems to partially confirmed our conjecture of the relationship between S4 subsite’s structure and cleavage efficiency: two residues within 3C^pro^ participating in forming backbone H-bond with substrate P4 position, N126 and S128, have same orientation but different coordinates in EV71 and CVA16.

Further study still need to be carried out in order to demonstrate if the above rules hold between SVA and FMDV, as well as other picornavirus. If the accomodation ability of bindng pocket is truly relevant to cleavage efficiency, taking FMDV as an example: SVA 3C^pro^ has a wider S4 pockets than FMDV, while P4 is a small amino acid residue in FMDV poly-protein cleavage site (Fig. 2A), so SVA may show higher cutting efficiency when applying to cut FMDV poly-protein. On the other hand, in FMDV poly-protein, the amino acids occupying subsites S1 and S1’ are usually glutamic acid/glutamine and glycine/threonine (Fig. 2A), those side chains size are similar to those in SVA. Therefore, although the S1, S1’ pockets are smaller in SVA 3C^pro^, it may not significantly influence the cleavage efficiency when cutting FMDV poly-protein.

In summary, the structural studies on 3C^pro^ could help to reveal the cleavage mechanism of 3C^pro^ in picornaviruses, and provide theoretical basis for the subsequent design of VLP vaccines.

## Materials and Methods

### Bioinformatics analysis

The phylogenetic analysis of picornaviral 3C^pro^ was carried out by MEGA 7 (Kumar et al., 2016). And the 3C^pro^’s cleavage specificities is analyzed by ClustalW (Larkin et al., 2007), Espript (Robert and Gouet, 2014) and *k*pLogo (Wu and Bartel, 2017). All sequences were derived from GenBank, accession numbers are as follows: Poliovirus (PV), CAA24465; Hepatitis A virus (HAV), AAA45466; Human TMEV-like cardiovirus (HTCV),ACB29695; Human Coxsackievirus A16 (CVA16), ACV33370; Coxsackievirus B3 (CVB3), AAA42931; Human rhinovirus A (HRV-A), ACK37367; Human rhinovirus B (HRV-B), BAA00168; Human rhinovirus C (HRV-C), ACZ67658; Enterovirus D68 (EV68), ABL61317; Human enterovirus 71 (EV71), ACY00662; Foot-and-mouth disease virus - type A (FMDV-A), AZS18886; Foot-and-mouth disease virus - type O (FMDV-O), AAG45408; Foot-and-mouth disease virus - type SAT2 (FMDV-SAT2), AAQ11227; Equine rhinitis B virus (ERBV), CAA65615; Porcine teschovirus (PTV), CAB40546; Bovine rhinitis A virus (BRAV), AKA20760; Porcine sapelovirus (PSV), AAM33242; Encephalomyocarditis virus (EMCV), AAA43037.

### Protein production

The original viral strain selected in this study was Seneca valley virus isolate SVV-001 (GenBank accession no. DQ641257). The gene for 3C^pro^ was amplified using the primers 3C-forward (5’-GGAATTCCATATGCAGCCGAACGTAGATATGG-3’) and 3C-reverse (5’-CCGCTCGAGTTACTGCATAGTGGCCAG-3’). The gene was then subcloned into pET-28a via NdeI and XhoI restriction sites, which generated a protein gene followed by an N-teriminal poly-histidine tag coding sequence. The construct was verified by DNA sequencing. The plasmids were then transformed into Escherichia coli BL21 (DE3) competent cells. The expression of 3C^pro^ was induced by 0.5 mM IPTG at 18 °C for 16-20 hr.

### Protein purification and crystallization

The expressed protein was initially purified in an ÄKTA Start system (GE Healthcare) by immobilized metal-affinity chromatography (IMAC) on a 5 ml HisTrap HP column (GE Healthcare). And further purified by gel filtration on a Hi Load Superdex 75 column (GE Healthcare). Peak fractions were concentrated to 10 mg/ml in 20 mM HEPES, pH 7.5, containing 150 mM NaCl. Crystals were obtained by sitting drop vapor diffusion experiments at 18 °C with commercial screening kits (HamptonReserch). Optimization of crystallization conditions was performed manually by hanging drop vapor diffusion experiments at 4 °C and 18 °C.

### Data collection and structure determination

Crystals were transferred to a 1:1 mix of reservoir solution and cryo-protectant solution and frozen in a stream of N_2_ gas at 100 K immediately prior to data collection. The data collection was performed at the Shanghai Synchrotron Radiation Facility (SSRF) using beamline BL17U at a wavelength of 1.5418 Å(Shanghai, China) (Wang et al., 2018; Wang et al., 2016). The collected intensities were indexed, integrated, corrected for absorption, scaled, and merged using the HKL2000 package (Otwinowski and Minor, 1997).X-ray diffraction data were processed with CCP4 program suite(Collaborative Computational Project, 1994). The crystals structures were determined by molecular-replacement method by using FMDV 3C^pro^ structure (Protein Data Bank code 2BHG) as the search model (Lebedev et al., 2008; McCoy, 2007). The initial model was obtained by Phaser MR, manual model adjustment and subsequently refinement were performed with COOT and Refmac5 respectively(Emsley and Cowtan, 2004).

## Reference

Belsham, G., and Bøtner, A. (2015). Use of recombinant capsid proteins in the development of a vaccine against the foot-and-mouth disease virus. Virus Adaptation and Treatment 7, 11–23.

Berger, A., and Schechter, I. (1970). Mapping the active site of papain with the aid of peptide substrates and inhibitors. Philos Trans R Soc Lond B Biol Sci 257, 249–264.

Birtley, J.R., Knox, S.R., Jaulent, A.M., Brick, P., Leatherbarrow, R.J., and Curry, S. (2005). Crystal structure of foot-and-mouth disease virus 3C protease. New insights into catalytic mechanism and cleavage specificity. J Biol Chem 280, 11520–11527.

Collaborative Computational Project, N. (1994). The CCP4 suite: programs for protein crystallography. Acta Crystallogr D Biol Crystallogr 50, 760–763.

Cui, S., Wang, J., Fan, T., Qin, B., Guo, L., Lei, X., Wang, J., Wang, M., and Jin, Q. (2011). Crystal structure of human enterovirus 71 3C protease. J Mol Biol 408, 449–461.

Curry, S., Roqué-Rosell, N., Zunszain, P.A., and Leatherbarrow, R.J. (2007). Foot-and-mouth disease virus 3C protease: recent structural and functional insights into an antiviral target. Int J Biochem Cell Biol 39, 1–6.

Emsley, P., and Cowtan, K. (2004). Coot: model-building tools for molecular graphics. Acta Crystallogr D Biol Crystallogr 60, 2126–2132.

Kumar, S., Stecher, G., and Tamura, K. (2016). MEGA7: Molecular Evolutionary Genetics Analysis Version 7.0 for Bigger Datasets. Mol Biol Evol 33, 1870–1874.

Larkin, M.A., Blackshields, G., Brown, N.P., Chenna, R., McGettigan, P.A., McWilliam, H., Valentin, F., Wallace, I.M., Wilm, A., Lopez, R., et al. (2007). Clustal W and Clustal X version 2.0. Bioinformatics 23, 2947–2948.

Lebedev, A.A., Vagin, A.A., and Murshudov, G.N. (2008). Model preparation in MOLREP and examples of model improvement using X-ray data. Acta Crystallogr D Biol Crystallogr 64, 33–39.

Leong, L.E., Walker, P.A., and Porter, A.G. (1993). Human rhinovirus-14 protease 3C (3Cpro) binds specifically to the 5’-noncoding region of the viral RNA. Evidence that 3Cpro has different domains for the RNA binding and proteolytic activities. J Biol Chem 268, 25735–25739.

Lu, G., Qi, J., Chen, Z., Xu, X., Gao, F., Lin, D., Qian, W., Liu, H., Jiang, H., Yan, J., et al. (2011). Enterovirus 71 and coxsackievirus A16 3C proteases: binding to rupintrivir and their substrates and anti-hand, foot, and mouth disease virus drug design. J Virol 85, 10319–10331.

Matthews, D.A., Smith, W.W., Ferre, R.A., Condon, B., Budahazi, G., Sisson, W., Villafranca, J.E., Janson, C.A., McElroy, H.E., Gribskov, C.L., et al. (1994). Structure of human rhinovirus 3C protease reveals a trypsin-like polypeptide fold, RNA-binding site, and means for cleaving precursor polyprotein. Cell 77, 761–771.

McCoy, A.J. (2007). Solving structures of protein complexes by molecular replacement with Phaser. Acta Crystallogr D Biol Crystallogr 63, 32–41.

Mosimann, S.C., Cherney, M.M., Sia, S., Plotch, S., and James, M.N. (1997). Refined X-ray crystallographic structure of the poliovirus 3C gene product. J Mol Biol 273, 1032–1047.

Otwinowski, Z., and Minor, W. (1997). Processing of X-ray diffraction data collected in oscillation mode. Methods Enzymol 276, 307–326.

Polacek, C., Gullberg, M., Li, J., and Belsham, G.J. (2013). Low levels of foot-and-mouth disease virus 3C protease expression are required to achieve optimal capsid protein expression and processing in mammalian cells. J Gen Virol 94, 1249–1258.

Porta, C., Xu, X., Loureiro, S., Paramasivam, S., Ren, J., Al-Khalil, T., Burman, A., Jackson, T., Belsham, G.J., Curry, S., et al. (2013). Efficient production of foot-and-mouth disease virus empty capsids in insect cells following down regulation of 3C protease activity. J Virol Methods 187, 406–412.

Robert, X., and Gouet, P. (2014). Deciphering key features in protein structures with the new ENDscript server. Nucleic Acids Res 42, W320–324.

Walker, P.A., Leong, L.E., and Porter, A.G. (1995). Sequence and structural determinants of the interaction between the 5’-noncoding region of picornavirus RNA and rhinovirus protease 3C. J Biol Chem 270, 14510–14516.

Wang, Q.-S., Zhang, K.-H., Cui, Y., Wang, Z.-J., Pan, Q.-Y., Liu, K., Sun, B., Zhou, H., Li, M.-J., Xu, Q., et al. (2018). Upgrade of macromolecular crystallography beamline BL17U1 at SSRF. Nuclear Science and Techniques 29, 68.

Wang, Z., Pan, Q., Yang, L., Zhou, H., Xu, C., Yu, F., Wang, Q., Huang, S., and He, J. (2016). Automatic crystal centring procedure at the SSRF macromolecular crystallography beamline. J Synchrotron Radiat 23, 1323–1332.

Wu, X., and Bartel, D.P. (2017). kpLogo: positional k-mer analysis reveals hidden specificity in biological sequences. Nucleic Acids Res 45, W534–W538.

Zhang, X., Zhu, Z., Yang, F., Cao, W., Tian, H., Zhang, K., Zheng, H., and Liu, X. (2018). Review of Seneca Valley Virus: A Call for Increased Surveillance and Research. Front Microbiol 9, 940.

Zunszain, P.A., Knox, S.R., Sweeney, T.R., Yang, J., Roque-Rosell, N., Belsham, G.J., Leatherbarrow, R.J., and Curry, S. (2010). Insights into cleavage specificity from the crystal structure of foot-and-mouth disease virus 3C protease complexed with a peptide substrate. J Mol Biol 395, 375–389.

